# Prostate-Specific Antigen Dynamics Predict Individual Responses to Intermittent Androgen Deprivation

**DOI:** 10.1101/624866

**Authors:** Renee Brady, John D. Nagy, Travis A. Gerke, Tian Zhang, Andrew Z. Wang, Jingsong Zhang, Robert A. Gatenby, Heiko Enderling

## Abstract

**Background:** Intermittent androgen deprivation therapy (IADT) is an attractive treatment approach for biochemically recurrent prostate cancer (PCa), whereby cycling treatment on and off can reduce cumulative dose, limit toxicities, and delay development of treatment resistance. To optimize treatment within the context of ongoing intratumoral evolution, underlying mechanisms of resistance and actionable biomarkers need to be identified.

**Methods:** We have developed a quantitative framework to simulate enrichment of prostate cancer stem cell (PCaSC) dynamics during treatment as a plausible mechanism of resistance evolution.

**Results:** Simulated dynamics of PCaSC and non-stem cancer cells demonstrate that stem cell proliferation patterns correlate with longitudinal serum prostate-specific antigen (PSA) measurements in 70 PCa patients undergoing multiple cycles of IADT. By learning the dynamics from each treatment cycle, individual model simulations predict evolution of resistance in the subsequent IADT cycle with a sensitivity and specificity of 57% and 94%, respectively and an overall accuracy of 90%. Additionally, we evaluated the potential benefit of docetaxel for IADT in biochemically recurrent PCa. Model simulations based on response dynamics from the first IADT cycle identify patients who would or would not benefit from concurrent docetaxel in subsequent cycles.

**Conclusion:** Our results demonstrate the feasibility and potential value of adaptive clinical trials guided by patient-specific mathematical models of intratumoral evolutionary dynamics continuously updated with each treatment cycle.

**Translational Relevance:** Compared to continuous androgen deprivation therapy, intermittent androgen deprivation (IADT) has been shown to reduce toxicity and delay time to progression in prostate cancer. While numerous mathematical models have been developed to study the response to both continuous and intermittent androgen deprivation, very few have identified actionable biomarkers of resistance and exploited them to predict how patients will or will not respond to subsequent treatment. Here, we identify prostate-specific antigen (PSA) dynamics as the first such biomarker. Mechanistic mathematical modeling of prostate cancer stem cell dynamics that dictate prostate-specific antigen serum levels predicts individual responses to IADT with 90% overall accuracy and can be used to develop patient-specific adaptive treatment protocols, and potentially identify patients that may benefit from concurrent chemotherapy. Model results demonstrate the feasibility and potential value of adaptive clinical trials guided by patient-specific mathematical models of intratumoral evolutionary dynamics continuously updated with each treatment cycle.

## Introduction

Prostate cancer (PCa) is the most common type of cancer in American men and the second leading cause of cancer mortality (*1*). Following surgery or radiation, the standard treatment for hormone-sensitive PCa is continuous androgen deprivation therapy (ADT) at the maximum tolerable dose (MTD) with or without continuous abiraterone acetate (AA) until the tumor becomes castration resistant (*2*). Importantly, advanced PCa is not curable because PCa routinely evolves resistance to all current treatment modalities. Continuous treatment approaches fail to consider the evolutionary dynamics of treatment response where competition, adaptation and selection between treatment sensitive and resistant cells contribute to therapy failure (*3*). In fact, continuous treatment, by maximally selecting for resistant phenotypes and eliminating other competing populations, may actually accelerate the emergence of resistant populations – a well-studied evolutionary phenomenon termed “competitive release”.

In part to address this issue, prior trials have used intermittent ADT (IADT) to reduce toxicity and delay time to progression (TTP). However, these trials were typically not designed with a detailed understanding of the underlying evolutionary dynamics. For example, a prospective Phase II trial of IADT for advanced PCa included an 8-month induction period in which the patients were treated at MTD prior to beginning intermittent therapy (*4*). We have previously postulated that only a small number of ADT-sensitive cells would typically remain after the induction period, thereby significantly reducing the potential of intermittent treatment to take advantage of the evolutionary dynamics (*3*).

Fully harnessing the potential of intermittent PCa therapy requires identifying ADT resistance mechanisms, predicting individual responses, and determining potentially highly patient-specific, clinically actionable triggers for pausing and resuming IADT cycles. Progress in integrated mathematical oncology may make such analysis possible. Many mathematical models based on a variety of plausible resistance mechanisms have been proposed to simulate IADT responses (*3, 5–13*). While these models can fit clinical data, they often rely on numerous model variables and parameters that in combination fail to adequately predict responses and outcomes for individual patients (*9*). We hypothesize that prostate cancer stem cells (PCaSCs) may be, at least in part, responsible for tumor heterogeneity and treatment failure due to their self-renewing, differentiating and quiescent nature (*14–16*). Simulating longitudinal prostate-specific antigen (PSA) levels in early IADT treatment cycles could help identify patient-specific PCaSC dynamics to computationally forecast individual disease dynamics and reliably predict IADT response or resistance in subsequent treatment cycles.

The first evidence of stem cells in the prostate was provided by Isaacs and Coffey (*17*), who used androgen cycling experiments in rodents to show that castration resulted in involution of the prostate, while restored androgen levels resulted in complete regeneration of the prostate. These findings demonstrated that normal prostate depends on androgens for maintenance. A small population of androgen-independent stem cells within the prostate epithelium divide to give rise to amplifying cells, which do not directly depend on androgen for their continuous maintenance but respond to androgens by generating androgen-dependent transit cells. Approximately 0.1% of cells in prostate tumors express the stem cell markers CD44^+^/α2β1^hi^/CD133^+^ (*18*), and mice as well as patients treated with ADT have significantly increased PCaSC populations (*19, 20*). This suggests that evolution of or selection for pre-existing androgen-independent PCaSCs contributes to treatment resistance.

The purpose of this study is to evaluate individual PSA dynamics in early IADT treatment cycles as a predictive marker of response or resistance in subsequent treatment cycles. We hypothesize that patient-specific PCaSC division patterns underlie the measurable longitudinal PSA dynamics, and that a mathematical model of PCaSCs can be trained to predict treatment responses on a per-patient basis. Here, we present an innovative framework to simulate and predict the dynamics of PCaSCs, androgen-dependent non-stem PCa cells (PCaCs), and blood PSA concentrations during IADT. Our mathematical model of PCaSC enrichment is calibrated and validated with longitudinal PSA measurements in individual patients to identify model dynamics that correlate with treatment resistance. The model’s predictive power to accurately forecast individual patients’ responses to IADT cycles is evaluated in an independent patient cohort. These analyses suggest that PCaSC and PSA dynamics may potentially be used to personalize IADT, maximize time to progression, and ultimately improve PCa outcomes. The calibrated and validated model is then used to generate testable hypotheses about patients that may benefit from concurrent chemotherapy.

## Materials and Methods

### IADT Clinical Trial data

The Bruchovsky prospective Phase II study trial was conducted in 109 men with biochemically recurrent prostate cancer (*21*). IADT consisted of 4 weeks of Androcur as lead-in therapy, followed by a combination of Lupron and Androcur, for a total of 36 weeks. Treatment was paused if PSA has normalized (< 4 *μ*g/L) at both 24 and 32 weeks, and resumed when PSA increased above 10 *μ*g/L. PSA was measured every four weeks. Patients whose PSA had not normalized after both 24 and 32 weeks of being on treatment were classified as resistant and taken off of the study. We analyzed the data of 79 patients who had completed more than one IADT cycle. One patient was omitted for inconsistent treatment, seven were omitted due to the development of metastasis and/or local progression, and one was omitted due to taking multiple medications throughout the trial, resulting in 70 patients included in the analysis (**Fig. S1**). To calibrate and assess the accuracy of our model, the data was divided into training *(n = 35, 27 responsive, 8 resistant)* and testing *(n = 35, 28 responsive, 7 resistant)* cohorts, respectively matched for clinical response to treatment.

### Mathematical Model of IADT Response

We developed a mathematical model of PCaSC (*S*), non-stem (differentiated) cells (*D*), and serum PSA concentration (*P*) (**Fig. S2**). PCaSCs divide with rate λ (day^−1^) to produce either a PCaSC and a non-stem PCa cell with rate *1-ps* (asymmetric division) or two PCaSCs at rate *p*_*s*_ (symmetric division) with negative feedback from differentiated cells (*15*). Differentiated cells exclusively produce PSA at rate ρ (*μ*g/L day^−1^), which decays at rate ϕ (day^−1^). Unlike androgen-independent PCaSCs, differentiated cells die in response to androgen removal at rate α (day^−1^) (*33*). IADT on and off cycles are described with parameter *T*_*x*_, where *T*_*x*_ = 1 when IADT is given and *T*_*x*_ = 0 during treatment holidays.

The coupled mathematical equations describing these interactions are shown below.

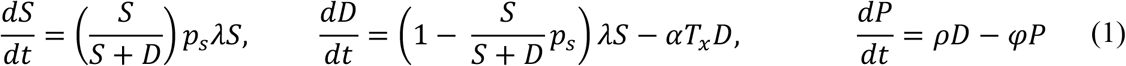

### Mathematical Model Training and Validation

We assume uninhibited PCaSCs divide on average once per day and set λ = ln(2). To reduce model complexity and prediction uncertainty, we assume PSA production rate (ρ) and decay rate (ϕ) to be uniform between patients. PCaSC self-renewal rate (*p*_*s*_) and differentiated cell ADT sensitivity (α) are correlated and assumed to be patient-specific. We used particle swarming optimization (PSO) (*34*) to identify population uniform and patient-specific model parameters that minimize the least squares error between model simulation and patient data in the training cohort. The trained mathematical model is assessed for accuracy in the validation cohort. The learned population uniform parameters are kept constant for all patients, and PSO is performed to find appropriate values for *p*_*s*_ and α to produce accurate data fits.

### Adaptive Bayesian Response Prediction

In order to predict the evolution of resistance, we started by fitting the model to each cycle of the training cohort data individually. That is, finding the optimal values of *p*_*s*_ and α, while allowing ϕ and ρ to remain fixed at the values previously found, to fit one cycle of data at a time (**Fig. S3A**). We then measured the relative change in *p*_*s*_ between cycles and used this to generate cumulative probability distributions as shown in **Fig. S3B**. Sampling from the 95% confidence interval around the exponential curve relating *p*_*s*_ to α in cycle *i+1* (**Fig. S3C**), we found a corresponding α_*i+1*_. This *p*_*s,i+1*_ and α_*i+1*_ were used to predict the response in cycle *i+1*. This process was repeated 1000 times to generate 1000 predictions. In line with the trial by Bruchovsky et al. (*21*), resistance was defined as PSA increasing during treatment and/or a PSA level above 4 *μ*g/L at both 24 and 32 weeks after the start of a cycle of treatment. Simulations that satisfied either of these conditions were classified as resistant. Thus, for each cycle prediction we obtained a probability of resistance (P(Ω) = number of resistance predictions out of 1000 simulations).

To determine whether to categorize a patient as responsive or resistant based on our predictions, we used the results from the training cohort data to find a cycle-specific threshold κ_i_ for each cycle (**Fig. S3D**). If P(Ω)> κ_*i*_, then the patient was predicted as resistant in cycle *i.* For each cycle, a value of κ was chosen that would maximize the sensitivity (predicting resistance when a patient is indeed resistant), and specificity (predicting resistance when a patient is actually responsive) of the training cohort. Each resistance threshold was used to predict response or resistance in the training cohort.

### Modeling concurrent docetaxel

Unlike ADT, docetaxel can induce cell death in both PCaSCs and non-stem cells, though to a lesser degree in PCaSCs compared to non-stem cells (*35*). To model this, we extended the current model to include death of each cell type at rates δ_*S*_ (day^−1^) and δ_*D*_ (day^−1^). That is,

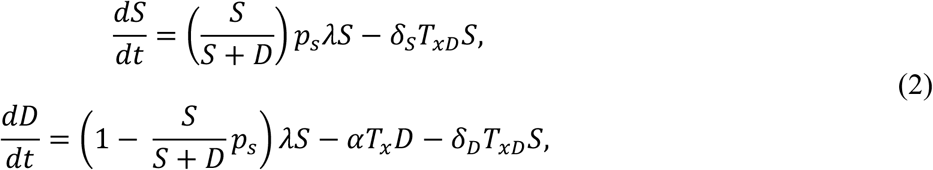

where *T*_*xD*_ = 1 when docetaxel is on and *T*_*xD*_ = 0 otherwise. Each cycle of docetaxel was simulated as a single dose on day *n* (*T*_*xD*_ = 1) followed by three weeks without docetaxel (*T*_*xD*_ = 0). The parameters δ_*S*_ = 0.0027 and δ_*D*_ = 0.008 were chosen such that approximately three times more non-stem cells died than PCaSCs (*35*).

## Results

### Mathematical model accurately simulates patient-specific IADT response dynamics

The model was calibrated to and assessed for accuracy on longitudinal data from a prospective Phase II study trial conducted in 109 men with biochemically recurrent prostate cancer treated with IADT (*21*) (see Materials and Methods, **Fig. S1**). Stratified random sampling was used to divide the data into training and testing cohorts. Assuming that uninhibited PCaSCs divide approximately once per day (*22*), λ (day^−1^) was set to ln(2) and parameter estimation was used to find the remaining parameters. The model was calibrated to the training cohort data with two population-uniform parameters (PSA production rate ρ = 1.87E-04 (*μ*g/L day^−^1), decay rate ϕ = 0.0856 (day^−1^)) and two patient-specific parameters (median PCaSC self-renewal rate *p*_*s*_ = 0.0278 [2.22E-14,0.2583] (non-dimensional), ADT cytotoxicity rate α = 0.0360 [0.0067,1] (day^−1^)). The model results captured clinically measured longitudinal PSA dynamics of individual responsive and resistant patients (**Fig. S4A, S4B**) and the population as a whole (R^2^ = 0.74 **Fig. S4C**). The corresponding PCaSC dynamics demonstrated a rapid increase in the PCaSC population in patients that became castration resistant compared to patients that remained sensitive throughout the trial. Model simulations also showed that emergence of resistance is a result of selection for the PCaSCs during on-treatment phases. Coinciding with this transition to castration resistance, analysis of model parameters revealed that resistant patients had significantly higher PCaSC self-renewal rates than responsive patients (median *p*_*s*_ = 0.0249 [1.18E-13,0.0882] for responsive patients vs. *p*_*s*_ = 0.0820 [2.2E-14,0.2583] for resistant patients, p < 0.001, **Fig. S4D**). As shown in **Fig. S4D**, analysis revealed an exponential relationship between *p*_*s*_ and α and therefore these parameters are not independent. This relationship allows prediction of future cycles based on PCaSC self-renewal rate *p*_*s*_ as the single, identifiable, independent patient-specific parameter.

To assess the accuracy of our model, we set the PSA production rate ρ and decay rate ϕ to the values obtained from the training cohort and identified patient-specific values for *p*_*s*_ and corresponding α in the testing cohort. With these, the model was able to fit the data equally well (R^2^=0.69) and the resulting parameter distributions and relationships were similar to those found in the training cohort (Fig. 1A-D).

**Fig. 1.**
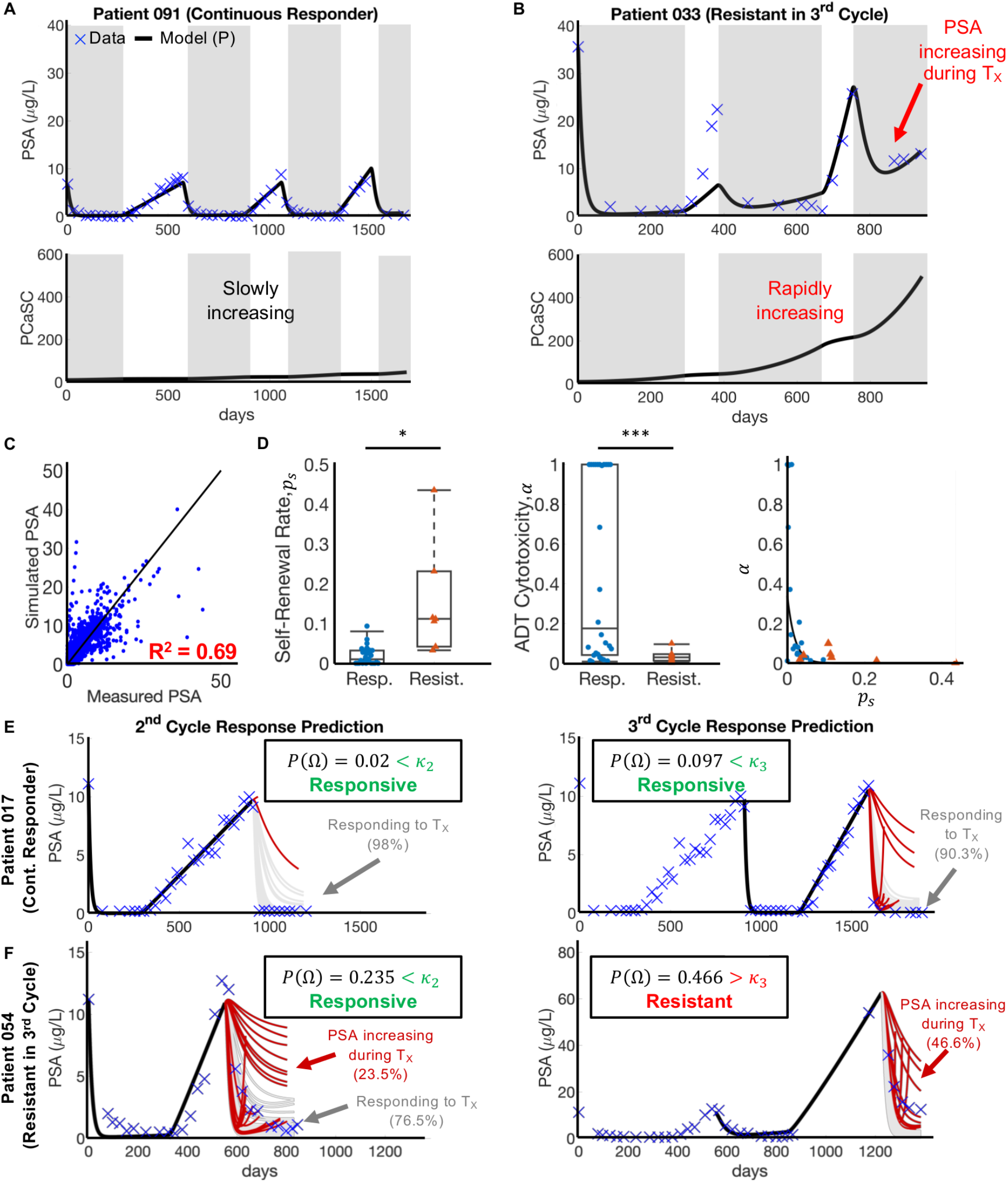
Model validation on testing patients. (**A** and **B**) Model fits to PSA data and corresponding PCaSC dynamics for (A) a continuous responder and (B) a patient who developed resistance during his third cycle of treatment. PCaSC population is rapidly increasing in resistant patient and slowly in responsive patient due a significantly higher self-renewal rate (*p*_*s*_ = 0.0201 and 0.1118 for patients 091 and 033, respectively). (**C**) Simulated vs. measured PSA. Linear regression obtains an R^2^ of 0.69. **(D)** Parameter distributions for the stem cell self-renewal *p*_*s*_ and ADT cytotoxicity *α*, with *ϕ* and *ρ* learned from training patients show similar trend found in training cohort. **(E)** The model predicted responsiveness in cycles two (98% of simulations) and three (90.3% of simulations) for Patient 017. The probability of resistant (*P*(Ω)) was less than its respective κ (κ_2_ = 0.45, κ_3_ = 0.29) for each cycle. He completed the trial on day = 2202. **(F)** The model predicted resistance in 23.5% of cycle 2 simulations and in 46.6% of cycle 3 simulations for Patient 054. The predicted probability of resistance in cycle three was greater than κ_3_ so he would be advised to stop the trial. Data showed that he became resistant on day = 1384 during the third cycle, as predicted.

### PCaSC self-renewal rates underlying observable PSA dynamics can predict subsequent IADT cycle responses

To predict the evolution of resistance in subsequent treatment cycles, we fit the model to single treatment cycles for patients in the training set, again setting the PSA production rate ρ and decay rate ϕ to the values previously found. The self-renewal and ADT cytotoxicity rates maintained the exponential relationship, previously obtained when optimizing over all IADT cycles. The distributions of the self-renewal rate *p*_*s*_ and the corresponding ADT cytotoxicity rate α, as well as their relative change from cycle to cycle, were used to predict responses in subsequent cycles of patients in the training cohort (see Materials and Methods). An appropriate resistance threshold was obtained for each cycle based on the forecast simulations of the training set. Using these thresholds, model forecasting was completed on the testing set and the patient was classified as either responsive or resistant in the subsequent treatment cycle. Representative examples of second and third cycle predictions for one responsive patient and one resistant patient from the testing cohort are shown in Fig. 1E-F. Patient 017 was a continuous responder who underwent three cycles of IADT before the end of the trial. The model correctly predicted responsiveness in cycles two and three based on the parameters fitted in cycles one and two, respectively. Patient 054 became resistant in the third IADT cycle and the model was able to correctly predict response in cycle two and resistance in the third cycle based on the thresholds learned in the training set. The model yielded a sensitivity of 57% and specificity of 94% over all subsequent IADT cycles for patients in the test cohort. The overall accuracy of the model was 90%.

### IADT without induction period significantly increases TTP

In a study by Crook et al., continuous ADT was compared to IADT in localized PCa patients and they found that IADT was non-inferior to ADT using overall survival as the clinical endpoint (*4*). A similar study by Hussain et al. conducted in metastatic, hormone sensitive PCa patients found that neither regimen proved superior (*23*). These findings are likely the result of the 7-8 months induction period, which resulted in the competitive release of the resistant phenotype once the androgen sensitive subpopulation was eliminated (*3*). Administering IADT without such an induction period would allow sufficient time for the sensitive subpopulation to efficiently compete with the resistant subpopulation and prolong time to progression and ultimately, overall survival.

To test these hypotheses, we simulated continuous ADT and IADT without the 36-week induction period and compared predicted TTP against the trial by Bruchovsky et al. (*21*) For IADT simulations, treatment was administered until PSA fell below 4 *μ*g/L and resumed again once it rose above 10 *μ*g/L at simulated measurements every 4 weeks. Both protocols were simulated until PSA progression or for 10 years (end of simulation, EOS), with progression defined as three sequential increases in PSA during treatment. For the IADT protocol, progression was also defined as PSA > 4 *μ*g/L at both 24 and 32 weeks on treatment, as used in the Bruchovsky trial. Comparing the TTP from the trial results against simulated continuous ADT showed that IADT with an induction period resulted in significantly longer TTP (Fig. 2A). Though median TTP was not reached for either IADT protocol, Fig. 2B shows that the average TTP was approximately 5 months longer for IADT without the induction period when compared to the actual trial data and the continuous ADT simulations. Fig. 2C shows TTP could be increased by more than 7 months in selected patients.

**Fig. 2.**
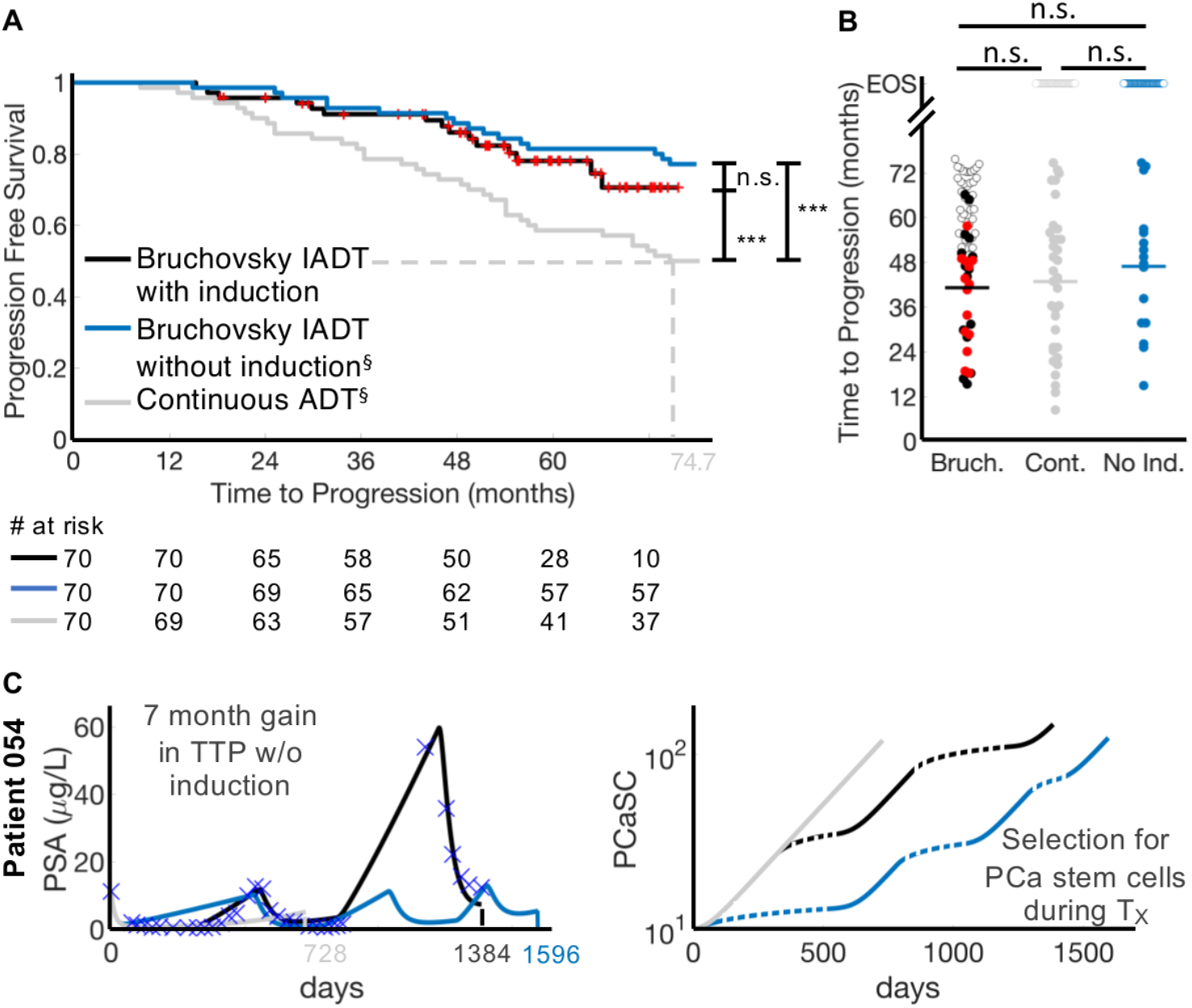
Administering IADT without induction period increases TTP when compared to IADT with induction and continuous ADT. **(A)** Kaplan-Meier estimates of progression comparing the Bruchovsky IADT protocol with induction (black curve) against simulated IADT without induction (blue) and continuous ADT (gray) (^§^ indicates predictive model simulation). With and without induction period, IADT TTP significantly increases when compared to continuous therapy TTP. **(B)** TTP comparison between Bruchovsky IADT with induction (black), continuous therapy (gray), and IADT without induction (blue). Bruchovsky patients who were lost to follow up are shown in red. Open circles denote end of simulation (EOS). Solid lines denote mean TTP (41.18, 42.83, and 46.9 months for Bruchovsky IADT with induction, continuous therapy, and Bruchovsky without induction, respectively). Though median TTP was not reached for either IADT protocol (A), the average TTP is longer with IADT without induction, compared to IADT with induction and continuous ADT. **(C)** Model fit to Patient 054 (Bruchovsky IADT protocol with induction, black curve), simulation without induction (blue curve), and simulation of continuous ADT (gray). On the Bruchovsky protocol, the patient became resistant after 1384 days. Simulating continuous ADT would result in progression after 728 days. Simulating IADT without the induction would increase TTP by 7 months. PCaSC dynamics show that treatment selects for PCaSC population, accelerating resistance development. Dashed line represent time when ADT is off.

### Alternative treatment decision thresholds can further increase TTP

In the ongoing pilot study (NCT02415621) of IADT in metastatic, castration resistant PCa patients, IADT is paused after a decline in PSA to below 50% of pre-treatment PSA levels and is resumed once PSA returns to the pre-treatment level. At the time of writing, of the 18 patients currently enrolled for more than 12 months, four have developed PSA and radiographic progression. The median time to progression has not been reached, but is at least 20 months greater than a contemporaneous cohort treated continuously with MTD until progression, as well as published cohorts. The treatment group has received an average cumulative dose of less than half that of standard of care treatment (*3*).

To test the validity of our model, we simulated IADT using a 50% threshold, as well as 10% and 70% and compared the results against those obtained in the Bruchovsky trial. Pausing treatment once PSA falls below 70% and 10% of the pre-treatment PSA, checking every two weeks, results in significantly longer TTP when compared to the Bruchovsky trial protocol (Fig. 3). Additionally, the average cumulative dose could be significantly reduced depending on the threshold used. With a 10% threshold, the average cumulative dose is ~50% of that of the Bruchovsky study. A 50% threshold results in ~22% of the cumulative dose used in the Bruchovsky study, while 70% results in ~20%. Controlling for the total time that each patient participated in the trial results in about 30%, 15%, and 13% of the comulative dose used in the Bruchovsky study for thresholds of 10%, 50%, and 70%, respectively (**Fig. S5**).

**Fig. 3.**
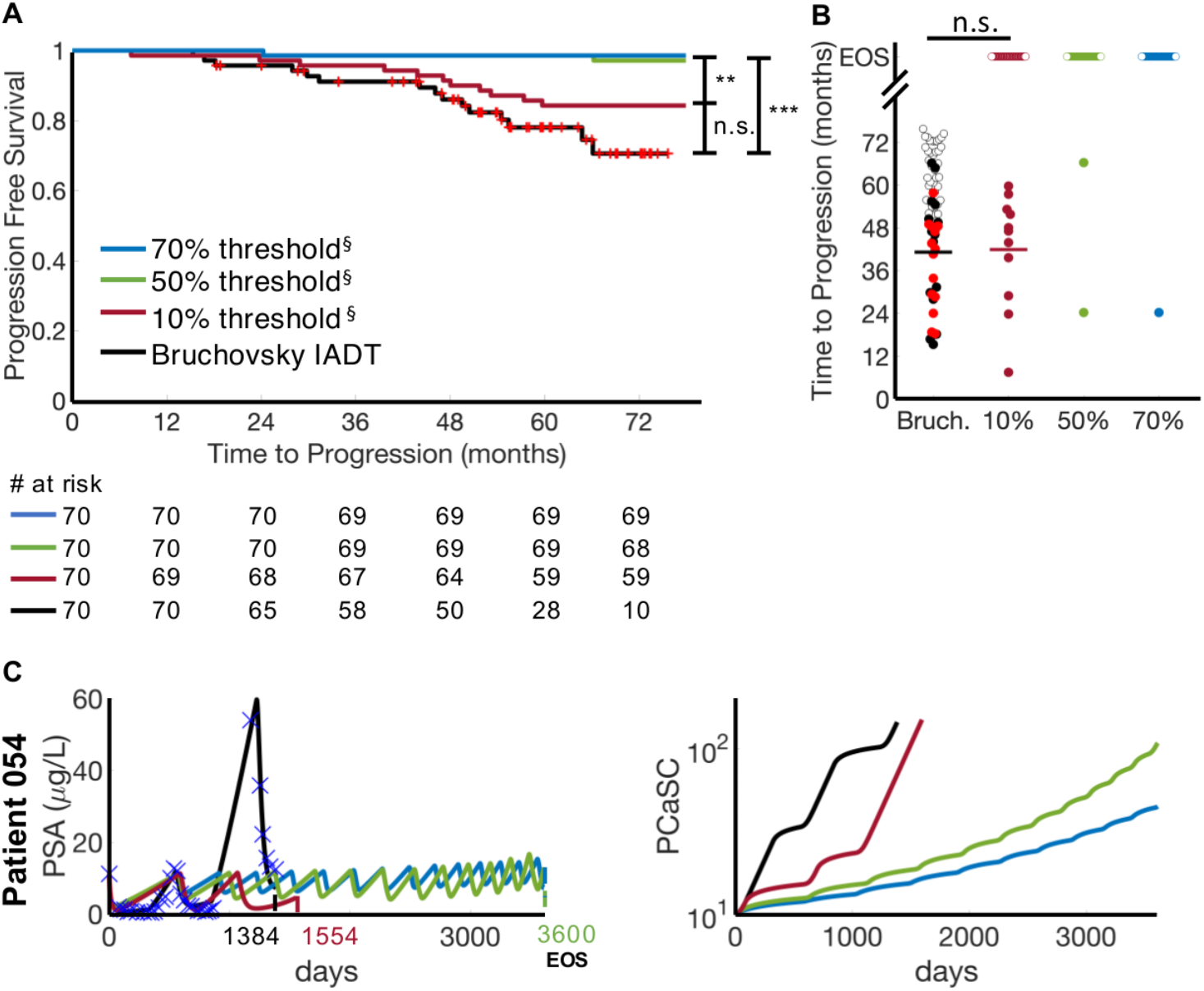
Alternative threshold therapy can significantly improve TTP. **(A)** Kaplan-Meier estimates of progression comparing the Bruchovsky IADT protocol (black curve) with alternative therapy protocol using a threshold of 10% (maroon), 50% (green), and 70% (blue) (^§^ indicates predictive model simulation). A 50% or 70% threshold can significantly increase TTP when compared to the Bruchovsky IADT protocol. **(B)** TTP comparison between Bruchovsky IADT (black) and thresholds of 10%, 50%, and 70%. Bruchovsky patients who were lost to follow up are shown in red. Open circles denote end of trial/simulation (EOS). Solid lines denote mean TTP (41.18 and 41.92 for Bruchovsky IADT and 10% trigger, respectively). **(C)** Alternative threshold therapy simulations for Patient 054. On the trial protocol, the patient became resistant after 1384 days. With a threshold of 10%, resistance could be delayed for 1554 days. With a threshold of 50% and 70%, the patient could continue on the protocol for more than 3600 days.

### Concurrent docetaxel administration provides favorable TTP

We sought to investigate the effect of docetaxel (DOC) in IADT in biochemically recurrent PCa patients by simulating six cycles of DOC with concurrent ADT, followed by IADT (as defined in the Materials and Methods section). Model analysis showed that patients with higher stem cell self-renewal rates would benefit the most from DOC in a castration naïve setting (prior to progression, Fig. 4A). Comparing progression free survival showed that castration naïve docetaxel could increase TTP (Fig. 4B). We also tested the ability of DOC to increase survival after the development of resistance to IADT. To do this, we simulated IADT, following the Bruchovsky trial protocol, and then added ten cycles of DOC plus continuous ADT after resistance developed. As shown in Fig. 4C, this increased mean TTP from 41.18 months observed in the Bruchovsky trial to model-predicted 50.14 months.

**Fig. 4.**
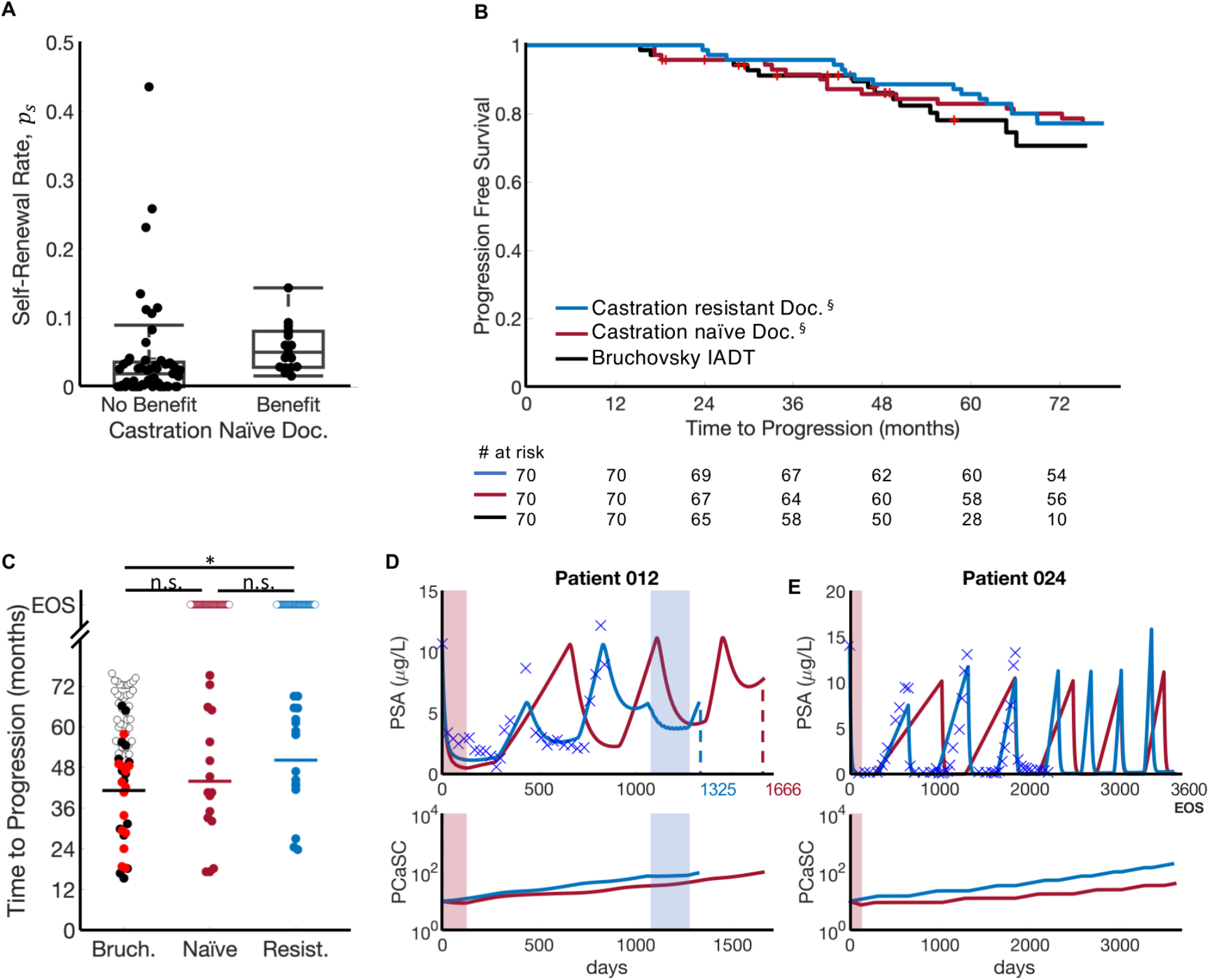
Docetaxel administration applied prior to or after IADT progression can increase TTP. **(A)** In those patients given docetaxel prior to IADT progression, patients with high *p*_*s*_ receive the highest benefit. **(B)** Kaplan-Meier estimates of progression comparing Bruchovsky IADT protocol (black) against castration naïve (maroon) and castration resistant (blue) docetaxel simulations (^§^ indicates predictive model simulation). Castration naïve and resistant refer to docetaxel given prior to IADT and after IADT progression, respectively. Six cycles of docetaxel prior to IADT increased TTP by 21.8 months on average. Ten cycles of docetaxel after IADT progression increased TTP by 17.5 months on average. **(C)** TTP comparison between Bruchovsky IADT (black) and castration naïve (maroon) and castration resistant (blue) docetaxel simulations. Bruchovsky patients who were lost to follow up are shown in red. Open circles denote end of trial/simulation (EOS). Administering docetaxel after IADT progression significantly increases TTP. Solid lines denote mean TTP (41.18, 43.92, and 50.14 months for Bruchovsky IADT, castration naïve, and castration resistant, respectively). Comparison of TTP shows that late DOC administration can significantly extend TTP when compared to IADT alone. **(D-E)** Castration naïve (red) and castration resistant (blue) simulations. **(D)** On the Bruchovsky IADT protocol, Patient 012 developed resistance after 839 days. Administering docetaxel prior to IADT delayed progression for an additional 27 months, while docetaxel given after progression allowed the patient to remain on treatment for an additional 16 months. **(E)** Though the trial concluded after 6.5 years, simulations shows that Patient 024 could have remained on the Bruchovsky IADT protocol for at least 10 years. Giving docetaxel prior to IADT produced similar results.

### First cycle PCaSC self-renewal rate stratifies patients who can benefit from docetaxel

As shown above, the stem cell self-renewal rate plays a significant role in IADT and could accurately predict a patient’s response in subsequent cycles after just the first cycle. With this, we used the first cycle *p*_*s*_ value to simulate six cycles of DOC with concurrent ADT followed by IADT and found that patients with *p*_*s*_ ≥ med(*p*_*s*_) could benefit from DOC after the first cycle of IADT (Fig. 5A). There was a significant difference in TTP between patients with a high *p*_*s*_ and those with a low *p*_*s*_ both when simulating IADT with and without DOC after the first cycle (Fig. 5B). Though the difference in TTP was not significantly higher with DOC given after the first cycle (Fig. 5C), median TTP increased from 35 to 41.5 months in those patients with *p*_*s*_ ≥ med(*p*_*s*_), as shown in Fig. 5B.

**Fig. 5.**
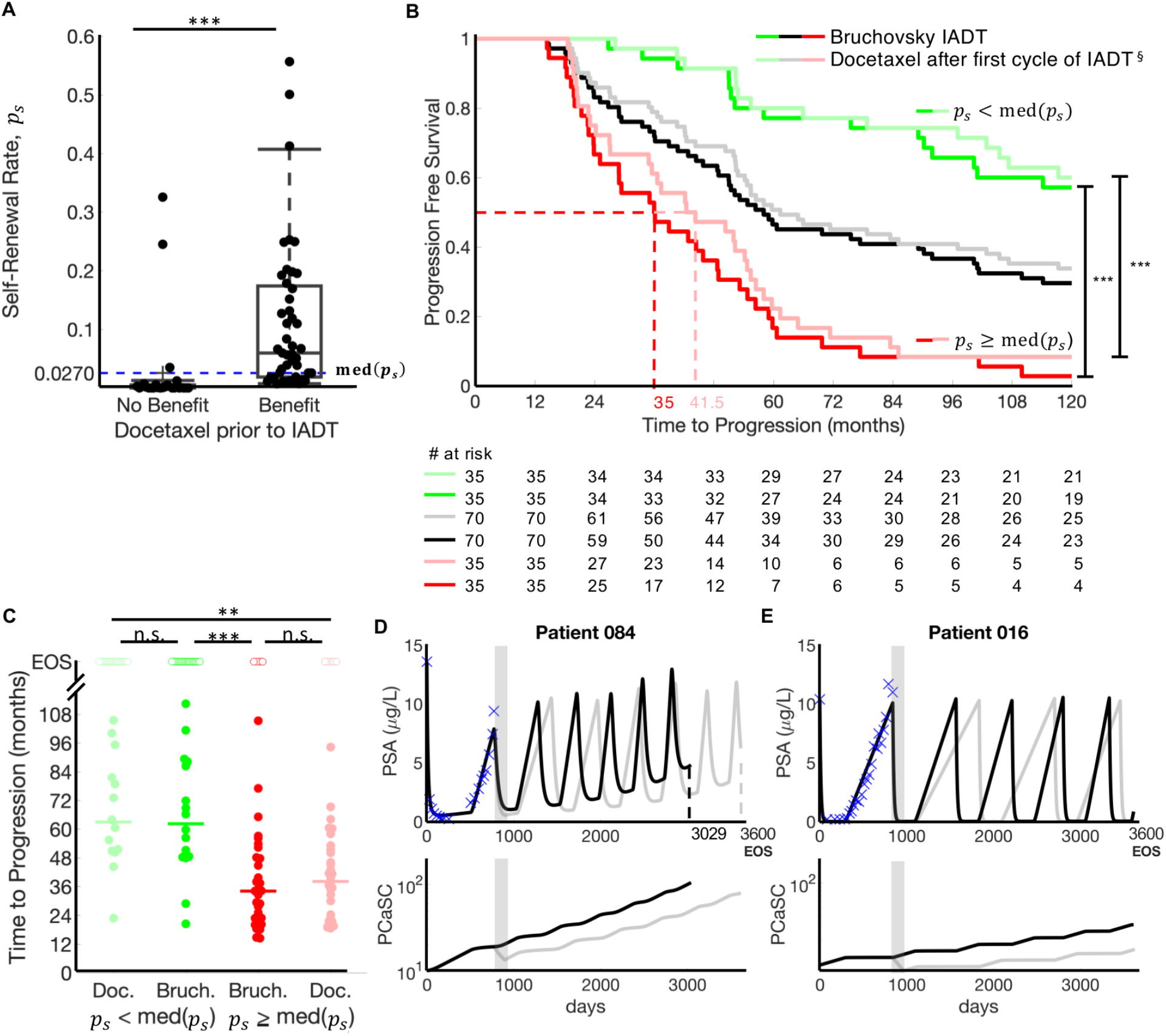
First cycle *p*_*s*_ stratifies patients who could benefit from docetaxel after first cycle of IADT. **(A)** *p*_*s*_ distributions between patients who may benefit from docetaxel after first cycle of IADT and those who may not. Median *p*_*s*_ stratifies patients who could benefit from docetaxel after first cycle of IADT. **(B)** Kaplan-Meier estimates of progression with and without docetaxel after first cycle of IADT (^§^indicates predictive model simulation). Stratifying by median *p*_*s*_ shows that patients with *p*_*s*_ ≥ med(*p*_*s*_) (red) experience greatest benefit from docetaxel (gain of 6 months). For patients with with *p*_*s*_ < med(*p*_*s*_) (green), there was not a significant benefit in TTP. **(C)** TTP comparison between Bruchovsky IADT with and without docetaxel. Open circles denote end of simulation (EOS). TTP was significantly lower in patients with *p*_*s*_ ≥ med(*p*_*s*_) both with and without docetaxel. Solid lines denote median TTP (62.98, 62.27, 34.13, and 38.15 months (left to right) for patients with low *p*_*s*_ (green) and high *p*_*s*_ (red), respectively). D-E. Simulation results of Bruchovsky IADT with (gray) and without (black) docetaxel after first cycle. **(D)** Patient 084 (*p*_*s*_ = 0.0386 > med(*p*_*s*_)) gains at least 18 months with docetaxel after first cycle. **(E)** Patient 016 (*p*_*s*_ = 0.0017 < med(*p*_*s*_)) did not progress with or without docetaxel.

## Discussion

Androgen deprivation therapy is not curative for advanced prostate cancer, as patients often develop resistance. IADT is a promising approach to counteract evolutionary dynamics by reducing competitive release of the resistant subpopulation during treatment holidays. Since IADT is highly dynamic, maximum efficacy requires continuous, accurate estimates of sensitive and resistant subpopulations.

Here we present a simple mathematical model of evolutionary dynamics within biochemically recurrent prostate cancer during IADT. The model has been trained with two parameters that are uniform across all patients and only two patient-specific parameters, which are interconnected, allowing us to further reduce them to a single, measurable parameter for each patient. Model simulations support the central hypothesis that the evolution of PCaSCs is highly correlated with the development of resistance to IADT. Resistant patients are likely to have higher PCaSC self-renewal rates than responsive patients, leading to increased production of PCaSCs and ultimately differentiated cells, thereby accelerating PSA dynamics with each treatment cycle. Our results are similar to prior studies in glioblastoma that cancer stem cell self-renewal is likely to increase during prolonged treatment (*24*).

Using longitudinal PSA measurements and observed clinical outcomes from the IADT trial by Bruchovsky et al., the model was calibrated to clinical data and predicted the development of resistance with a 90% accuracy. The theoretical study produced three important clinical findings: (1) IADT outcomes in prior studies were adversely affected by the 8-month induction period, which reduced IADT overall survival comparable to that of continuous ADT. (2) With IADT, applying a PSA treatment threshold that depends on pre-treatment PSA levels (rather than a fixed value for all patients) can significantly increase TTP. (3) Early treatment response dynamics during IADT can identify patients that may potentially benefit from concurrent docetaxel treatment, particularly patients with a large stem cell self-renewal rate after one cycle of IADT.

Our study also demonstrated the value of ongoing model simulations in predicting outcomes from each treatment cycle throughout the course of therapy. By accumulating data from each cycle to continuously estimate the current tumor population dynamics, model simulations could predict the response to the next cycle with a sensitivity and specificity of 57% and 94%, respectively and an overall accuracy of 90%. This ability to learn from prior treatments and predict future outcomes adds an important degree of flexibility to a cancer treatment protocol– a game theoretic strategy termed “Bellman’s Principle of Optimality” that greatly increases the physician’s advantage (*25*).

For those predicted to become resistant in the next cycle of IADT, an ideal model would also predict alternative treatments that could produce better clinical outcomes. The role of docetaxel in metastatic, hormone-sensitive PCa has been investigated in three studies in the past five years. The GETUG-AFU15 study found a non-significant 20% increase in overall survival in high volume disease (HVD) patients who received DOC concurrently with continuous ADT, but no survival benefit in those with low volume disease (LVD) (*26*). Subsequently, the results of the STAMPEDE trial found that DOC administration resulted in a more than 12 months overall survival benefit with a median follow-up of 43 months (*27*). Finally, the CHAARTED trial showed a statistically significant overall survival benefit from adding DOC in patients with HVD; however, no statistically significant survival benefit was found in LVD patients (*28*). Here we explored the option of adding docetaxel in the treatment of biochemically recurrent PCa. We found that estimating the PCaSC self-renewal rate *p*_*s*_ using data from the first cycle of IADT could stratify patients who would receive the most benefit from concurrent administration of docetaxel. These results emphasize the critical heterogeneity within patients that affect response to therapy and the important role of quantitative models in identifying patient-specific parameters and defining appropriate treatment protocols based on model predictions.

Our study has some limitations in both the clinical data set and quantitative models. Since only 21% of the 70 patients progressed before the conclusion of the trial (*21*) (**Fig. S1**), the clinical data included more responsive than resistant patients in both the training and testing cohorts. Additionally, the majority of patients progressed within their second or third cycle, with only 24 patients entering the fourth cycle. The scarcity of available data during the fourth cycle made finding an appropriate resistance threshold challenging. A larger training data set would increase the sensitivity and overall accuracy of the model.

A limitation of the quantitative model is the use of dynamic PSA values as the sole biomarker of PCa progression. PCa can become aggressive and metastatic despite low levels of serum PSA (*29, 30*) with development of androgen independent prostate cancer, most notably neuroendocrine prostate cancer. Additional serum biomarkers such as circulating tumor cells (CTCs) and cell-free DNA (cfDNA) may prove useful in estimating intratumoral evolutionary dynamics in subsequent trials. With the CellSearch platform, higher CTCs enumeration >5cells per 7.5mL of peripheral blood has been shown to be prognostic and portend worse overall survival in metastatic CRPC patients (*31*); however detecting CTCs in biochemically recurrent patients has been labor-intensive with low yield (*32*).

In conclusion, our study demonstrates that a simple mathematical model based on cellular dynamics in prostate cancer can have a high predictive power in a retrospective data set from patients with biochemically recurrent PCa undergoing IADT. In particular, we demonstrate the model can use data from each treatment cycle to estimate intratumoral subpopulations and accurately predict the outcome of subsequent cycles. Furthermore, in patients who are predicted to fail therapy in the next cycle, alternative treatments for which a response is more likely can be predicted. We conclude that PSA dynamics can prospectively predict treatment response to IADT, suggesting ways to adapt treatment to delay TTP.

## Acknowledgements

We thank participants of the clinical trial and Dr. Bruchovsky for sharing the data. We would also like to thank our PSOC patient advocate Mr. Robert Butler for fruitful discussions.

## Author contributions

A.Z.W., T.Z., J.D.N. and H.E. conceptualized the study. R.B, J.D.N, T. A. G., and H.E. performed the modeling and statistical analyses. R.B, J.D.N., T.A.G., T.Z., A.Z.W., R.A.G, and H.E. wrote the manuscript.

