## Supplementary Figures for "Prostate-Specific Antigen Dynamics Predict Individual Responses to Intermittent Androgen Deprivation"

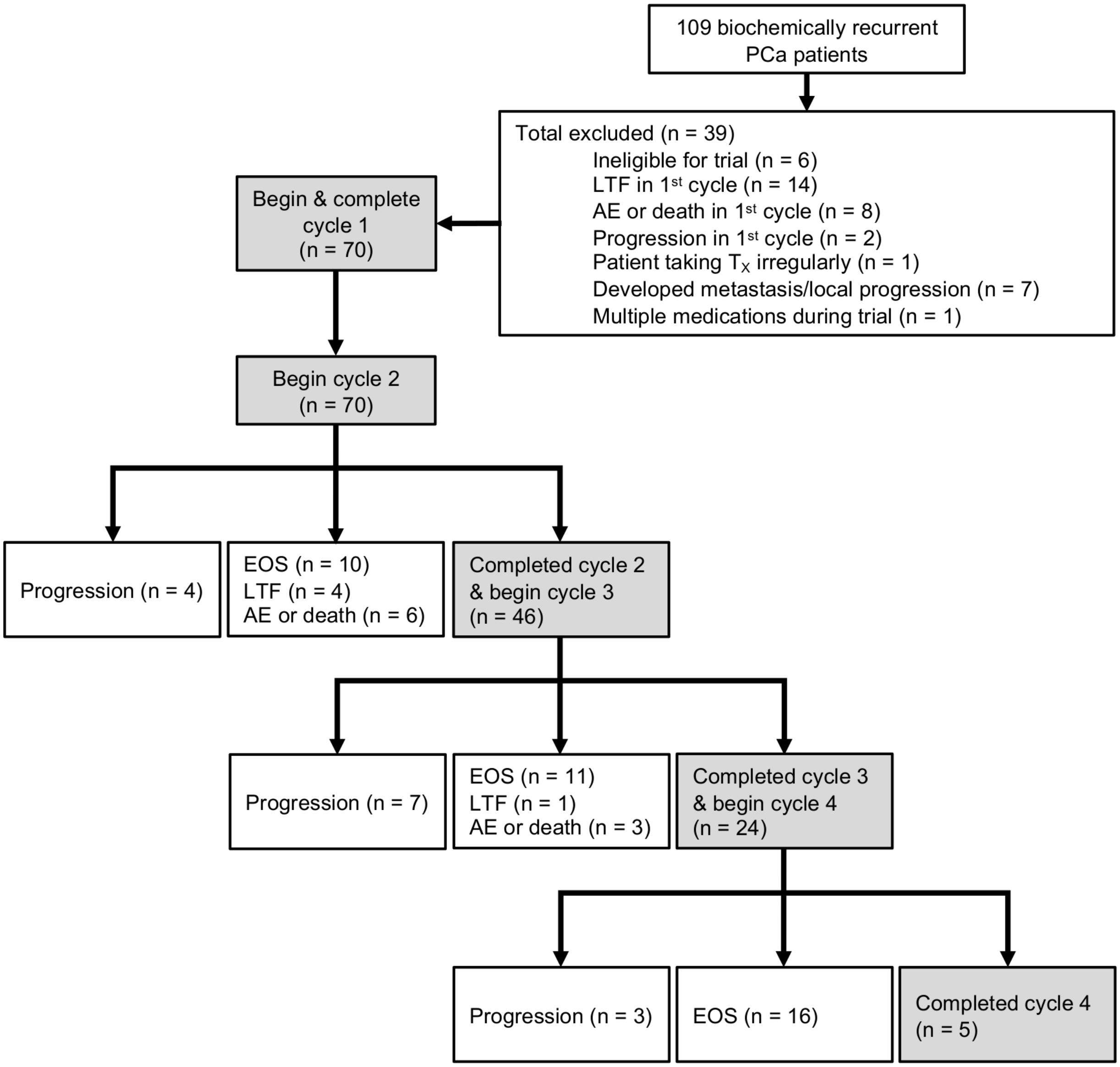


**Fig. S1. Data stratification for biochemically recurrent PCa patients enrolled in trial by Bruchovsky et al.** Of the 109 patients enrolled in the trial, 70 were included in the analysis. EOS, LTF, and AE denote end of study, lost to follow up, and adverse event, respectively.

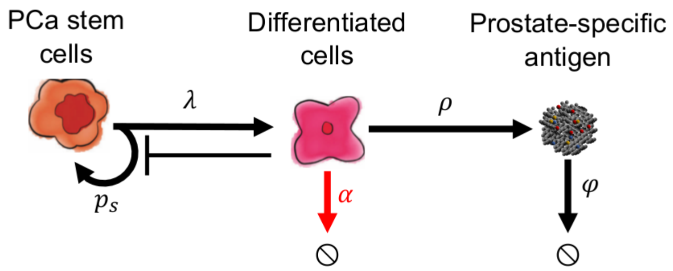


**Fig. S2. Model of prostate cancer stem cell and non-stem (differentiated) cell dynamics.** Interactions between stem and non-stem cells, as well as serum PSA. Prostate cancer stem cells can divide asymmetrically to produce differentiated cells. The differentiated cells inhibit the production of PCaSCs, die in response to ADT (shown by the red arrow), and produce PSA. Parameters are stem cell proliferation rate ($\lambda$), stem cell self-renewal $p_s$, ADT cytotoxicity $\alpha$, PSA production rate $\rho$, and PSA decay rate $\varphi$.

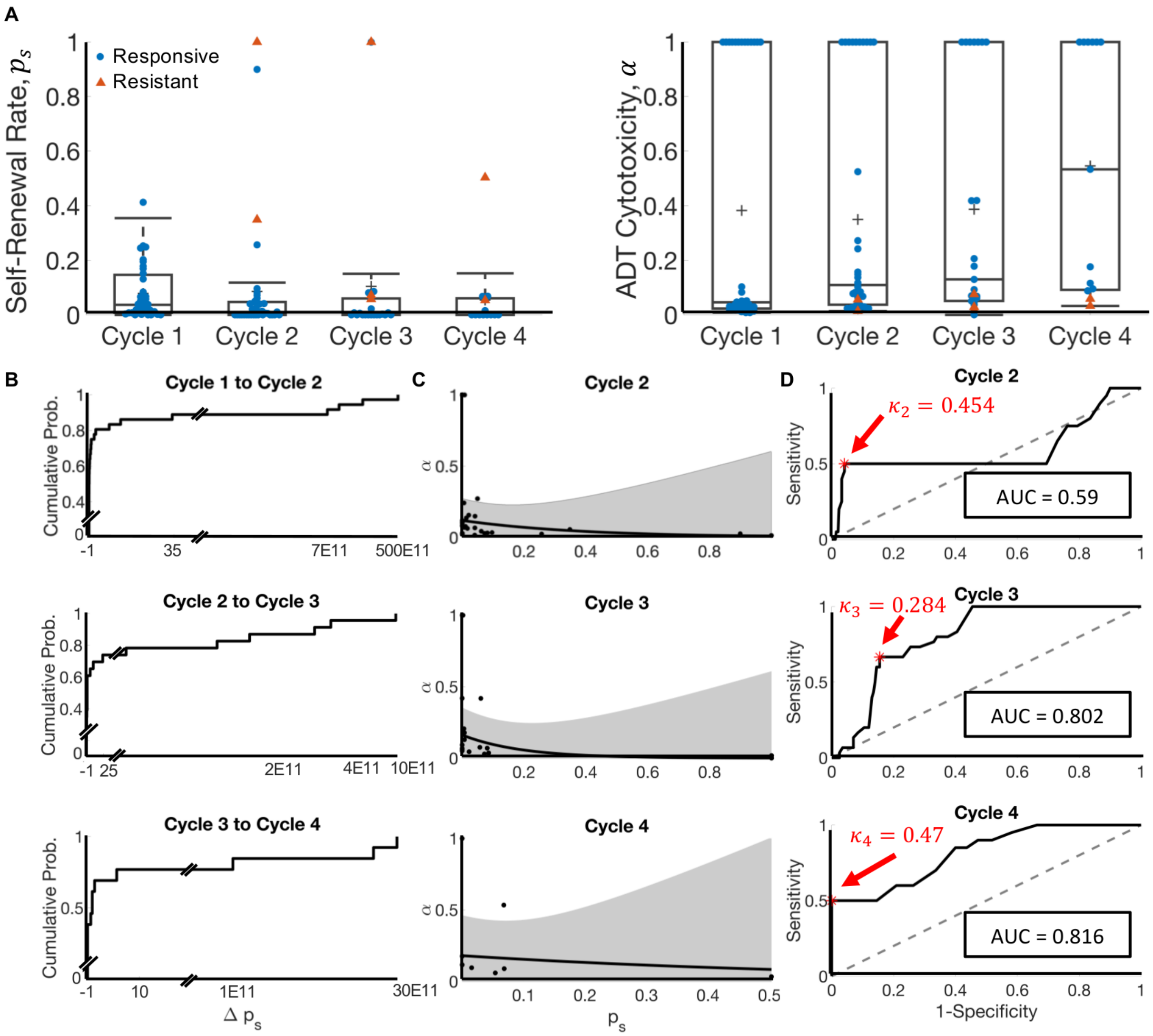


**Fig. S3. Cycle to cycle parameter changes. (A)** Cycle to cycle parameter distributions for $p_s$ and $\alpha$ (changes between cycles is not significant (p > 0.05)**. (B)** Cumulative probability of relative changes in $p_s$ between cycles. **(C)** $\alpha$ vs. $p_s$, fit (black curve), and 95% confidence interval (gray). **(D)** Receiver operating curves for training predictions for cycles two through four. The resistance threshold, $\kappa$, that maximizes the accuracy and hence, predictive power, is shown in red.

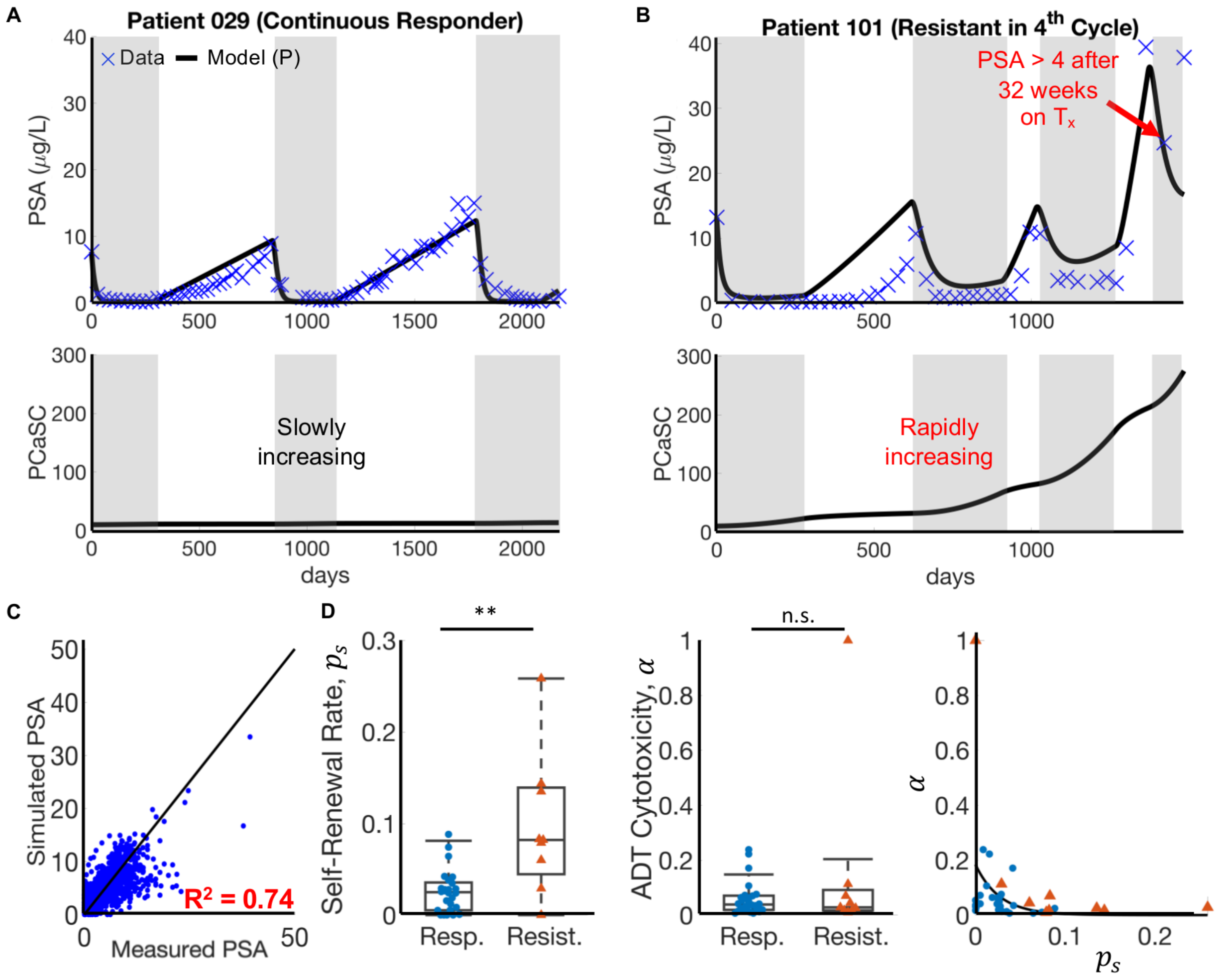


**Fig. S4. Parameter distributions and model fits for training patients.** (**A** and **B**) Model fits to PSA data and corresponding PCaSC dynamics for (A) a continuous responder and (B) a patient who developed resistance during his fourth cycle of treatment. PCaSC population is rapidly increasing in resistant patient and slowly in responsive patient due a significantly higher self-renewal rate ($p_s$ = 0.0052 and 0.1349 for patients 029 and 101, respectively) **(C)** Simulated vs. measured PSA. Linear regression obtains an $R^2$ of 0.74. **(D)** Parameter distributions, with $\varphi$ and $\rho$ uniform between all training patients. Stem cell self-renewal $p_s$ and ADT cytotoxicity $\alpha$ exhibit exponential relationship.

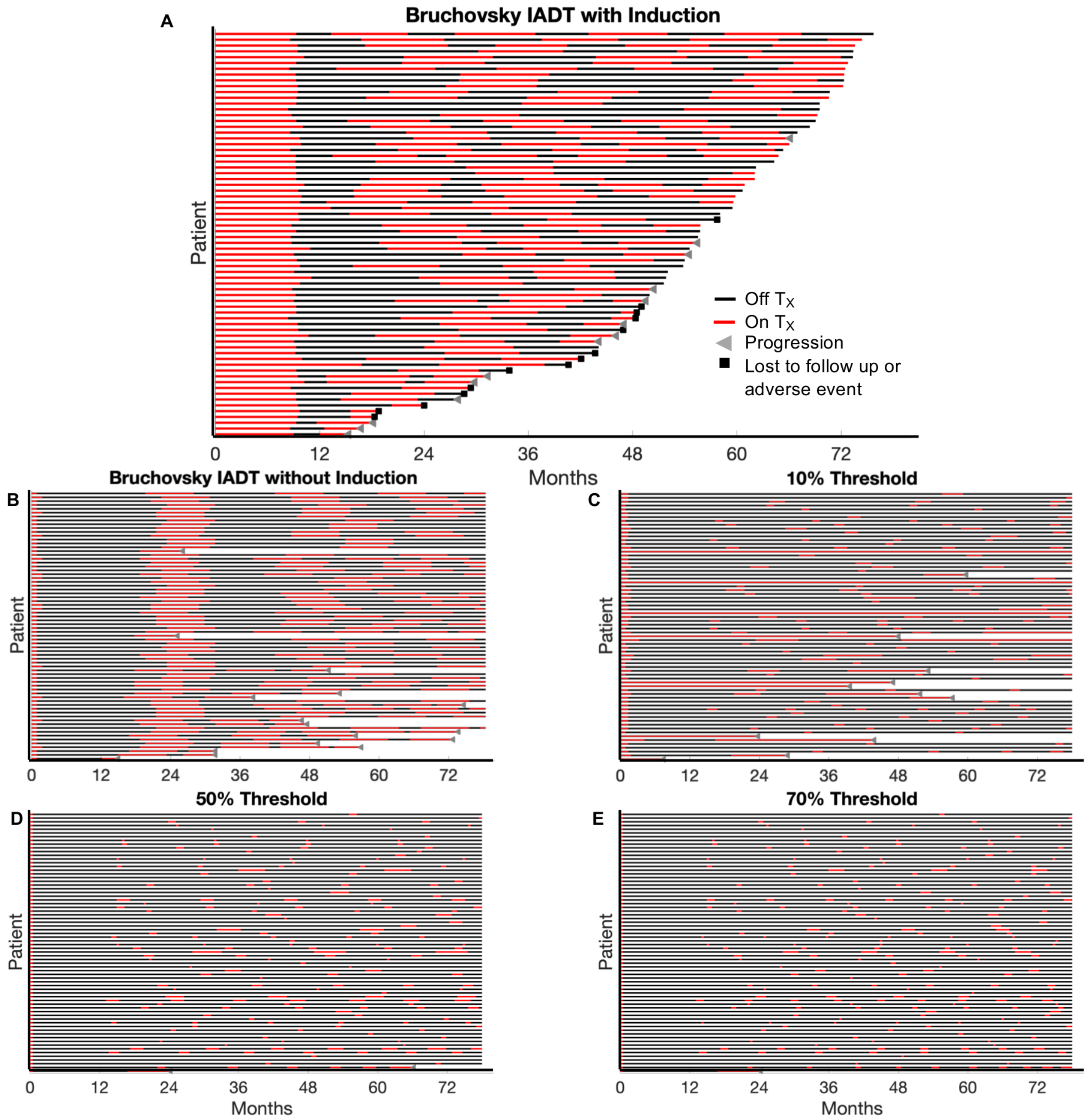


**Fig. S5. IADT treatment timing for Bruchovsky patients and simulated IADT without induction and with varying thresholds. (A)**Treatment times for IADT patients from trial by Bruchovsky et al. (with induction period). Average time on $T_x$/cycle = 8.96 months, average total time on $T_X$ = 27.65 months. **(B)** Simulated treatment times for IADT without induction period. Average time on $T_x$ /cycle = 6.88 months, average total time on $T_X$ = 24.6 months. **(C)** Simulated treatment times for IADT using a PSA threshold of 10%. Average time on $T_x$ /cycle = 4.51 months , average total time on $T_X$ = 13.13 months. **(D)** Simulated treatment times for IADT using a PSA threshold of 50%. Average time on $T_x$ /cycle = 1.24 months, average total time on $T_X$ = 6.01 months**. (E)** Simulated treatment times for IADT using a PSA threshold of 70%. Average time on $T_x$ /cycle = 0.94 months, average total time on $T_X$ = 5.1 months.
